# *In-silico* comparative modeling and interaction studies of PRSV proteins - Accelerated towards dissection of *structure based* evolutionary divergence and functional interaction of virus within the host

**DOI:** 10.1101/2022.09.09.507246

**Authors:** Anam Saleem, Zahid Ali, Saadia Naseem

## Abstract

Function based structure analysis of viral proteins reinforce their distinctive and unanticipated role within the host. The interaction dynamics of virus protein and host which is prerequisite for complete infectivity as well as systemic spread of invading virus demands to explore viral proteins in framework of their interacting partners. *Papaya ringspot virus* strain from Pakistan (PRSV-PK) spreading as atypical variant and causes drastic reduction in papaya production. The in-depth knowledge of the virus variant and effective management is obligatory. The desired objective is achievable once the evolutionary dynamics, molecular characterization and physicochemical structural properties conceived from the 3D protein structures are comprehended. Although the diversity studies on PRSV-PK strain been established but still there is a niche regarding structural based evolutionary dynamics of virus proteins and their probable interaction mode inside the host. The present investigations provided insights into the *in-silico* analysis of the functionally significant genes Coat protein (CP), Helper component proteinase (HC-Pro) and Nuclear Inclusion b protein (NIb) of PRSV-PK. The protein structure has been modeled using Phyre2, Swiss-Model and i-TASSER. Phyre2 built model showed 100% confidence for 67%, 63% and 20% sequence identity residues for PRSV CP, HC-Pro and NIb proteins respectively. The Swiss model showed identity values of 63.40%, 62.42% and 16.49% for CP, HC-Pro and NIb protein. whereas, i-TASSER server exhibited identity values of 67%, 63% and 19% for CP, HC-Pro and NIb proteins respectively. These structures provided a base line for functional analysis of experimentally derived crystal structures. The predicted models were validated using protein structure checking tools PROCHECK. Further the PRSV-PK-CP structures were compared with the PRSV-CP structures of representative isolates from different geographical regions. Nevertheless, the comparative modeling provides the insight into the evolutionary characteristics and proposed genetic diversity of PRSV based on protein structures. In addition, the conserved functional motifs have been mapped on aligned protein sequences of CP, HC-Pro and NIb, and their critical function within the host has been highlighted. Eventually, the interaction of papaya protein with the invading PRSV-CP has been predicted through *in-silico* protein-protein docking to elucidate their possible role in virus inhibition. The established structural-functional relation provided a basis to propose probable host-virus interactions in terms of virus infectivity, resultant host adaptability, and host defense response activation to counteract the virus invasion.

## Introduction

The evolutionary interaction between plant and pathogens in terms of their endeavors for survival and domination plays a foremost role in agriculture success [1]. Plants are well adapted to resist the attack of invading biotic and abiotic stresses. However, the biotic agents are of major concern in food production, where $30 billion annually of crop losses are attributed to viral diseases [2]. The highly adaptive nature of viruses and the numerous molecular modes they adopt enables them to take over the host cellular machinery to their benefit. At the same time, they must also avoid host defense responses activated in response to virus invasion, to ensure complete infectivity [3]. Thus, the evolving strategies of the invading virus and the resultant host defense elicitation can be fully understood only when the evolutionary dynamics of virus at nucleotide as well as protein structural level are fully explored. Perhaps the structural studies of invading viral proteins would provide better insight to the geographical strain divergence at protein level, a foundation to explore host-virus interactions in terms of virus infectivity and the resultant host adaptability eliciting either its own defense response or else the host defense triggered in response to invading virus. The 3D structure prediction using amino acid sequence are striking tools introduces where advanced AlphaFold is a powerful AI system developed by DeepMind to predict 3D structure of a protein from its amino acid sequence [4].

In this study we have worked on *Papaya ringspot virus* (PRSV) which is the predominant plant virus transmitted by aphids to *Cucurbitaceae* and *Caricaceae* families [5]. A part from the global papaya destruction, atypical strain of PRSV has severely affected papaya grown in Pakistan [6,7]. The nucleotide based molecular and evolutionary studies provide sufficient insight into the genetic variability of the virus strain [8–12]. However, the differential role of essential virus proteins in terms of virus divergence and evolution could also be clarified through their structural prediction [13]. The genome of PRSV contains different proteins including: Helper component Proteinase. PI (Protease), P3, CI, Nuclear inclusion and Nuclear inclusion b proteins.

The interaction between viral protein and host chromatins plays a significant part in viral infections and this interaction further modulates the host chromatin. Viral proteins of potyviruses family, including RNA-dependent RNA polymerase, nuclear inclusion NIa, NIb, CI helicase, HC-Pro, and P3 protein are all translocated in nucleus and interact differently with the host cellular proteins. There are several reports describing the interaction of NIa protein of *Turnip mosaic virus* (TuMV) with the poly (A) binding protein as well with the initiation factor eIF(iso) 4E in the nucleus. Thus, invading plant viruses modulate and utilize the host chromatin machinery and make their infection process more efficient. Research is also being carried to bridge a gap to understand the interaction between invading viral proteins and host nucleosome [14].

The potyviral coat protein (CP) is indulged in various functions during the infectious cycle, as viral RNA translation, replication, movement and encapsidation [15]. Moreover, the CP has remained an authentic target gene for developing PRSV resistant papaya and in this regard the nucleotide sequence based evolutionary divergence has already been predicted [6]. However, the comparative structural mapping of functionally active residues of CP would not only augment the previously reported viral divergence but also provide insight into the functional interaction required for virus invasion, movement and defense suppression inside the host. Another essential, HC-Pro (50-53 KDa) protein of potyviruses has been known to indulge in various functions of virus life cycle including the helping role of it in aphid based plant-plant transmission [16]. Proteolysis of viral poly protein [17,18], genome amplification and infectivity, virus movement in the host, RNA binding interaction with host proteins and most importantly interference of plant defense via gene silencing suppression are other functions of this protein. Thus, the structural aspects of HC-Pro, addressed in this study are helpful in reinterpretation of its functions in plant regulatory mechanisms. Another universal viral protein encoded by all RNA viruses is the RdRp (replicase protein) which enables to analyze the evolutionary history of RNA viruses. This gene namely the NIb gene is located between the NIa-Pro cistron and the CP-coding sequence in most of the potyviruses genomes [19] and is absolute requirement for genome replication. The structural analysis of these replicase proteins suggest the position of conserved domains and motifs and the functional role associated with them [19]. The structural features of the viral proteins in one way or other are definitely involved in tuning the host adaptability, and activation of host defense in response to viral infection. Henceforth, this study has been designed to predict the structural significance of essential PRSV-PK proteins and also the significant differences these protein possess, compared to their aggressive counterparts at distant geographical locations.

In view of papaya crop importance in industry and its potential to boost up economy in Asia-Pacific region, it has been mandatory to explore the structural based divergence of invading virus based on its essential proteins. The elaborated structural comparative homology modeling of the targeted full length CP, HC-Pro and NIb proteins would provide divergence information and possibly the way that how interaction of the viral protein with host factors play role in plant defense. For analysis and protein structures prediction of the respective genes modeling was done by using most topical homology modeling tools including Swiss-Model, Phyre2 and i-TASSER.

## Materials and Methods

### Sequence retrieval and analysis

The complete gene sequences of Coat protein (CP), Helper component proteinase gene (HC-Pro) and Nuclear inclusion b protein (NIb) of PRSV-PK with GenBank Accession numbers were retrieved from NCBI (http://www.ncbi.nlm.nih.gov/) as available on October, 30 2021.

### Protein Modeling of PRSV-PK genes

The nucleotide sequences of CP, HC-Pro and NIb were translated into amino acid sequences using EXpasy Translate software [20]. The physicochemical characterization of amino acid sequences of three genes was carried out by Expasy’s ProtParam server (http://web.expasy.org/protparam/). ProtParam (References / Documentation) a tool that allows the computation of various physical and chemical parameters for a given protein stored in Swiss- Prot or TrEMBL or for a user entered protein sequence. The parameters including the molecular weight, theoretical isoelectric point (pI), amino acid composition, atomic composition, extinction coefficient, estimated half-life, instability index, aliphatic index and grand average of hydropathicity (GRAVY) (Disclaimer) were computed. SOPMA server [21] (http://npsapbil.ibcp.fr/) was used for secondary structure prediction [22,23]. Further, the secondary structures were predicted using SOPMA (Self-optimized prediction method and alignment). Homology modeling of Coat protein, HC-Pro and NIb was performed using Phyre2, Swiss-Model and i-TASSER web servers. Information from amino acid sequence and homologous protein (with known structure) were used by the server to predict three-dimensional structure of protein [24].

### Phyre2 (Protein Fold Recognition Server)

Phyre2 is an automated homology modeling program with the increased alignment performances of a new alignment strategy using/rely on profiles or hidden Markov models (HMMs). The translated sequences of CP, HC-Pro and NIb of PRSV-PK were submitted to predict the protein model using Phyre2 Server.

#### Swiss-Model

Swiss-Model can be accessed via ExPASy web server, or from the DeepView (Swiss PDB-Viewer) program. An automated SWISS-MODEL [25] was used for three-dimensional (3D) protein structure modelling in this study. The interface of which is World Wide Web which provides several levels of interaction to users. Only amino acid sequences of available PRSV-CP, HC-Pro and NIb genes were used to build 3D model in “First Approach mode” whereas selection of template, alignment and modelling are performed by the server. The second is “Alignment mode” where target-template alignment is provided by user and modelling process is performed by server.

#### i-TASSER (Iterative threading ASSEmbly refinement)

i-TASSER [26,27] is a hierarchical protocol for automated protein structure prediction and also performs structure-based function annotation. Using the amino acid sequence of target proteins, I-TASSER first generates full-length atomic structural models from multiple threading alignments and iterative structural assembly simulations followed by atomic-level structure refinement. The biological functions of the protein, including ligand-binding sites, enzyme commission number, and gene ontology terms, are then inferred from known protein function databases based on sequence and structure profile comparisons [26,27]. Translated sequences of three genes, CP, HC-Pro and NIb of PRSV were uploaded respectively to the software queue. Figure 1 shows the flow chart in which three automated server tools were used to generate protein structures of three significant proteins of PRSV.

**Figure 1.**
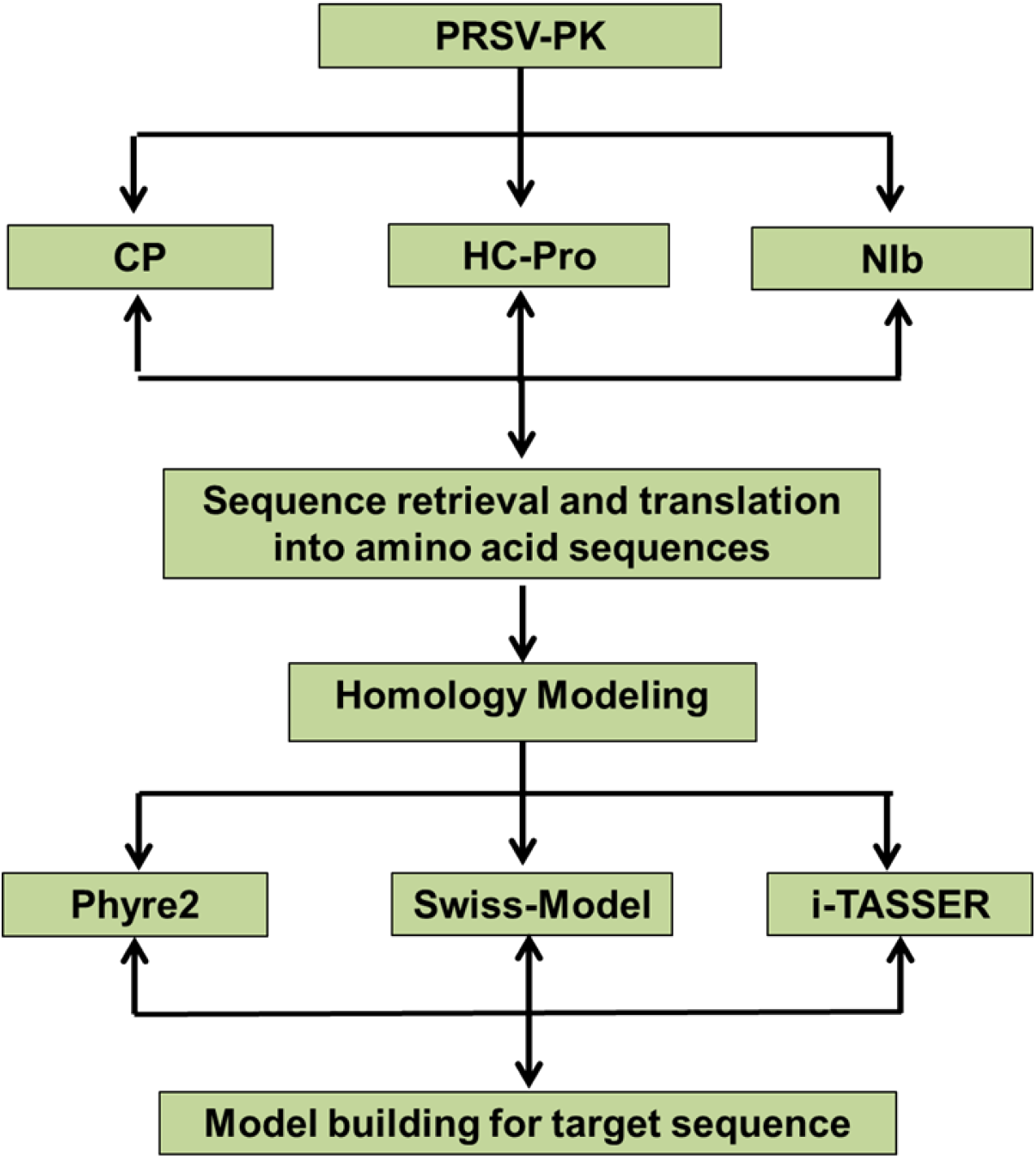
The flow chart showing step wise homology modeling of PRSV-PK proteins

### MODEL Validation

All the protein models of CP, HC-Pro and NIb proteins were evaluated using PROCHECK tool [28]. The PDB files of the build 3D structures were uploaded in PROCHECK tool. Ramachandran plot calculated in PROCHECK were used to validate the model quality based on the percentage residues lies in the favoured and disallowed regions which eventually confirmed the reliability of model stereochemistry.

### Structural Alignment of PRSV-PK-CP

The Coat protein structure of PRSV-PK was compared with the Coat protein structures of PRSV isolates from 3 representative clades (identified in section of genetic variability and evolutionary dynamics). Three isolates were selected from each clade i.e American clade (S46722_Hawaii, AJ012650_Mexico, KT275938_Colombia), South Asian clade (ABO44342_ Malaysia, AY010721_Thailand, X97251_Taiwan) and closely related clade including (MH397222_Bangladesh, MF356497_Meghalaya-India, LC482263_West Bengal-India) were selected. All the CP models of the representative isolates from each clade were developed through i-TASSER. The .pdb file of the structural models was superimposed with the PRSV-PK-CP structure using PYMOl2. The differences in the PRSV-PK amino acid residues with respect to other isolates were identified.

### *In-silico* Docking analysis of PRSV-PK CP with host proteins

A set of 11 crystal structures of wild type proteins belonging to *Caricaceae* were retrieved from RCSB-PDB [29] as well as from UNIPROT having an X-Ray resolution better than 3 Aº. The proteins were individually docked to PRSV-CP using the program Cluspro server, (https://cluspro.org) a widely used tool for protein-protein docking. The parameters such as cluster size, energy values and electrostatic values have been utilized for scoring the docking. Out of 10 possible docking modes suggested by program, the top ranking complexes were used for further analysis.

## Results

### Physicochemical properties of amino acids of CP, HC-Pro and NIb

The physicochemical characterization of CP, HC-Pro and NIb proteins of PRSV-P was done and the computation of various physical and chemical parameters for a given proteins are mentioned in Table 1. Grand average hydropathy (GRAVY) values of all the amino acids divided by the number of residues in the sequence, for the proteins CP. HC-Pro and NIb were -0.842, -0.435 and -0.328.

**Table 1.** Physicochemical characterization of Coat protein, HC-Pro and NIb by Expasy’s Protparam tool

| Sr. No | Physicochemical parameters | Coat Protein CP | Helper Component Proteinase HC-Pro | Nuclear Inclusion Protein Nib |
| --- | --- | --- | --- | --- |
| 1 | M.Wt (Da) | 32124.15 | 51800.94 | 61758.45 |
| 2 | PI | 7.77 | 7.07 | 5.49 |
| 3 | -R | 41 | 56 | 78 |
| 4 | +R | 42 | 55 | 62 |
| 5 | EC | 29910 | 49110 | 111325 |
| 6 | II | 34.52 | 36.51 | 37.32 |
| 7 | AI | 62.98 | 80.66 | 85.0 |
| 8 | GRAVY | -0.842 | -0.435 | -0.328 |

pI: Isoelectricpoint, +R: Positively charged residues, -R: Negatively charged residues, EC: Extinction coefficient, II: Instability Index, AI: Aliphatic Index, GRAVY: Grand Average Hydropathy Values.

### SOPMA (Self Optimized Prediction method and alignment)

The secondary structural features of three proteins were predicted using SOPMA (Self Optimized Prediction method and alignment). The findings revealed the domination of random coil in case of CP while domination of Alpha helix followed by other structure elements (Table 2).

**Table 2.** Physicochemical characterization of Coat protein, HC-Pro and NIb by SOPMA

| Sr. No | Parameters | CP | HC-Pro | NIb |
| --- | --- | --- | --- | --- |
| 1 | Alpha helix (Hh) | 39.72% | 43.54% | 52.97% |
| 2 | 3<sub>10</sub> helix (Gg) | 0.00% | 0.00% | 0.00% |
| 3 | Pi helix (Ii) | 0.00% | 0.00% | 0.00% |
| 4 | Beta bridge (Bb) | 0.00% | 0.00% | 0.00% |
| 5 | Extended Strand (Ee) | 10.28% | 14.88% | 13.01% |
| 6 | Beta Turn (Tt) | 4.26% | 5.47% | 5.95% |
| 7 | Bend region (ss) | 0.00% | 0.00% | 0.00% |
| 8 | Random coil (cc) | 45.74% | 36.11% | 28.07% |
| 9 | Ambiguous states | 0.00% | 0.00% | 0.00% |
| 10 | Other states | 0.00% | 0.00% | 0.00% |

### Homology Modeling of PRSV-PK proteins

#### Phyre2

The Models prediction of CP, HC-Pro, and NIb through Phyre2 generates protein models. The confidence percent and the % identity of the top models of three genes are mentioned in Table 3. The PDB structures of three genes generated are shown in the Fig. 2a (i, ii, iii). The Phyre2 server showed 100% confidence for sequence identity of 67%, 63% and 20% for CP, HC-Pro and NIb.

**Table 3.**
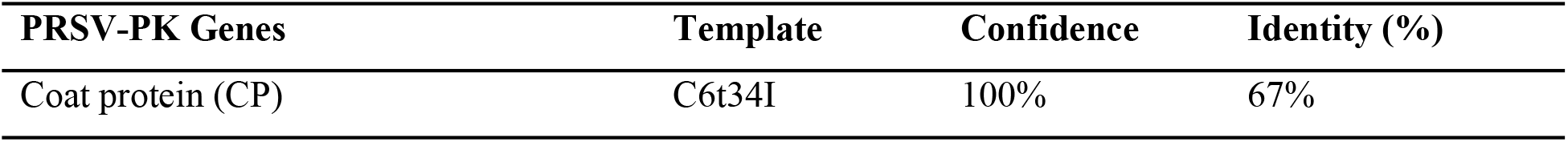
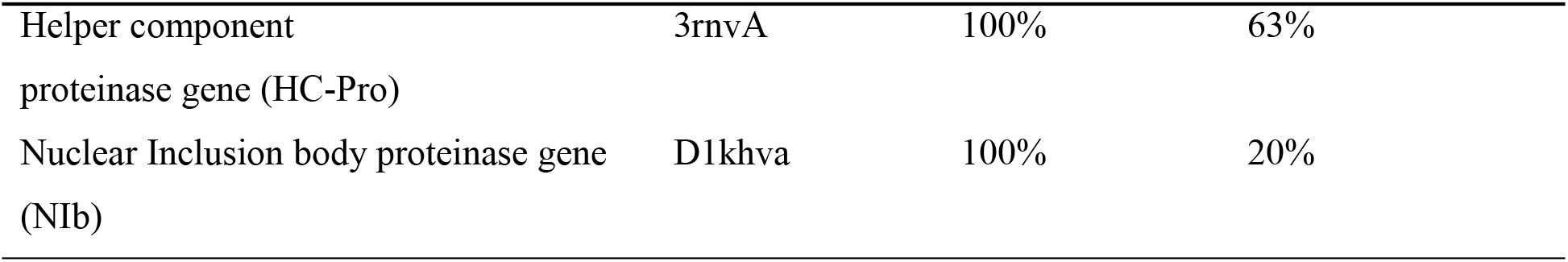
The confidence % and sequence identity of CP, HC-Pro and NIB genes retrieved through Phyre2

**Figure 2.**
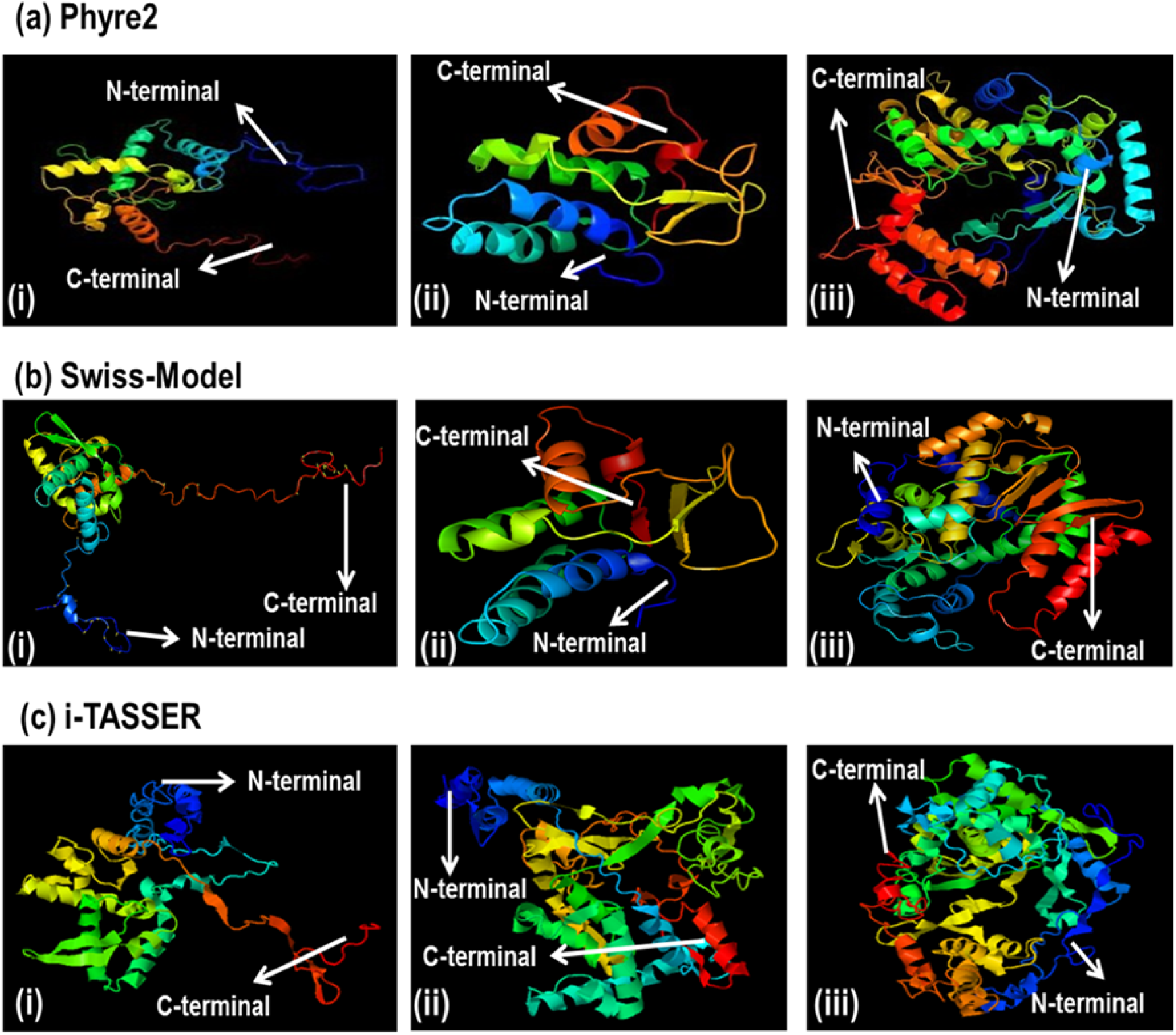
Three dimensional Models of PRSV-functional proteins built via homology modeling tools (a) Phyre2 built models of (i) Coat Protein (ii) HC-Pro (iii) NIb. (b) Models built by Swiss-Model (i) Coat Protein (ii) HC-Pro (iii) NIb (c) Models built by i-TASSER (i) Coat Protein (ii) HC-Pro (iii) NIb. The blue color in each structure represents the N-terminal region and Red color in each structure represents C-terminal region.

#### Swiss Model

The individual parameters for each gene calculated by the online Swiss modeler including QMean, Cß, All atoms, Solvation, Torsion, Template and sequence identity are mentioned in the Table 4. The predicted protein structures of three genes are shown in the Fig 2b (i, ii, iii).

**Table 4.**
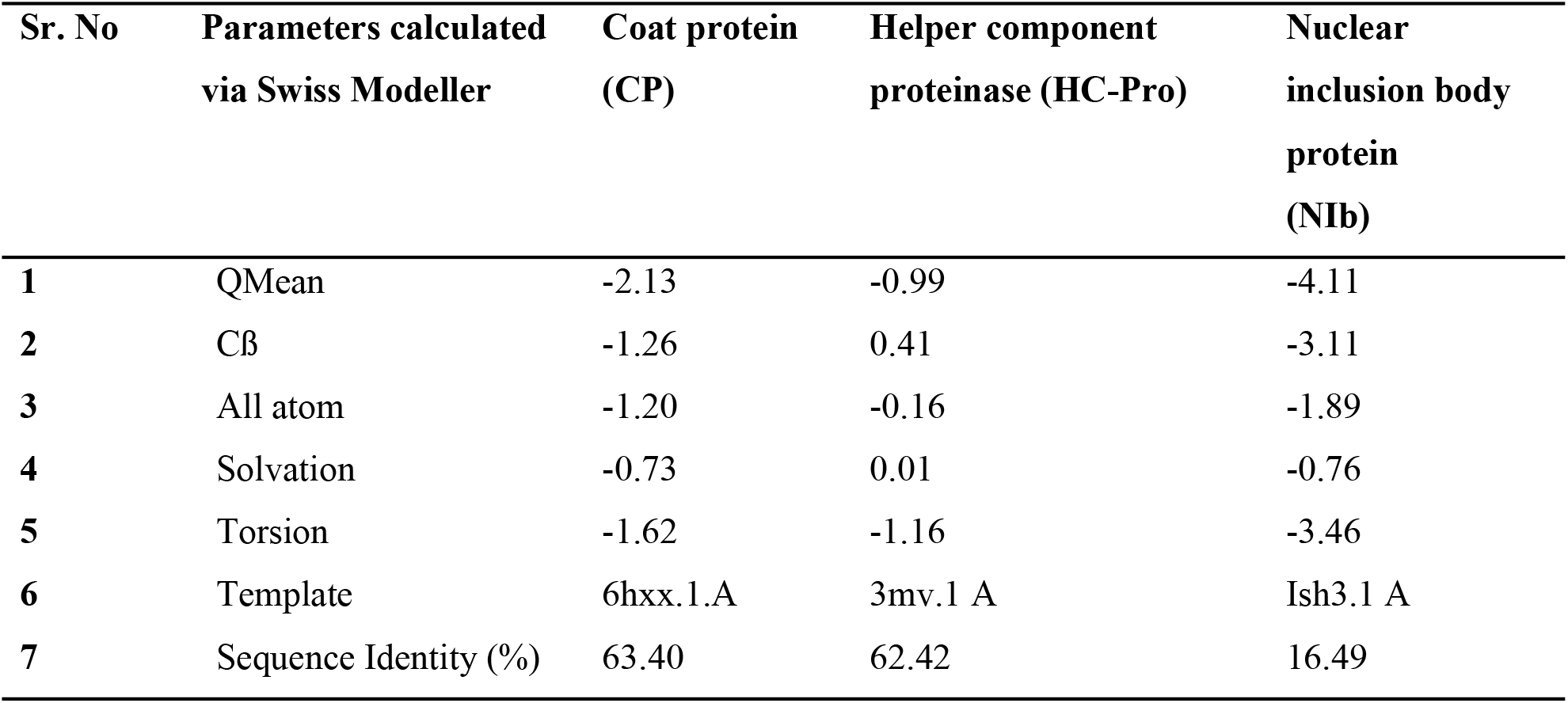
Parameters of predicted protein models of CP, HC-Pro and NIb calculated by Swiss-Modeler

#### i-TASSER Analysis

i-TASSER, an automated comparative protein structure modeling recognized the templates for one or more sequences based on the best available Protein Data Bank (PDB) template structures. The parameters of modeling computed for each individual protein model are mentioned in Table 5 and Figure 2c (i, ii, iii).

**Table 5.** The model template, c-score, T-score, estimated RMSD, ligand and ligand binding sites of CP, HC-Pro and NIB genes retrieved through i-TASSER

| PRSV-PK Protein Models | Template | Identity % | C-Score | Estimated T-score | Estimated RMSD | Ligand | Ligand binding sites |
| --- | --- | --- | --- | --- | --- | --- | --- |
| Coat protein (CP) | 6t34A | 67% | -2.27 | $0.44 \pm 0.14$ | $11.5 \pm 4.5\text{\AA}^{\circ}$ | Nuc Acid | 98, 99, 139, 141, 142, 143, 171, 173, 200, 217, 233, 237, 241 |
| Helper component proteinase (HC-Pro) | 3rnvA | 63% | -1.22 | $0.56 \pm 0.15$ | $9.9 \pm 4.6\text{\AA}^{\circ}$ | FES<br>DI-MU-SULFIDO-DIRON | 203, 204, 207, 211, 212, 213, 215 |
| Nuclear inclusion protein b (NIB) | 1khvA | 19% | -1.24 | $0.56 \pm 0.15$ | $10.4 \pm 4.6\text{\AA}^{\circ}$ | Nuc Acid | 124, 125, 130, 135, 170, 172, 188, 189, 190, 191, 200, 211, 221, 222, 223, 224, 309, 310, 311, 312, 314, 349, 431, 434, 438 |

### Model validation via PROCHECK

Ramachandran plot calculation in PROCHECK validation package measured the quality of the modeled structure built by Phyre2, Swiss Model and i-TASSER. The protein model for PRSV-PK HC-Pro built via Phyre2 showed highest percentage of residues (89.2%) residues in the allowed region and (0.0%) in the disallowed region showing the model is of good quality. The NIb model built by Swiss-Mod indicates (88.4%) residues in the allowed region and (1.2%) in the disallowed region. The Percent residues in allowed and disallowed regions are mentioned in Table 6.

**Table 6.** Validation of models built using PROCHECK tool

| Proteins | Model 1 (Phyre2) |  | Model 2 (Swiss-Model) |  | Model 3 (i-TASSER) |  |
| --- | --- | --- | --- | --- | --- | --- |
|  | Percentage of Residues in most favored region | Percentage of Residues in disallowed region | Percentage of Residues in most favored region | Percentage of Residues in disallowed region | Percentage of Residues in most favored region | Percentage of Residues in disallowed region |
| **PK-CP** | 75.4% | 0.5% | 87.5% | 0.5% | 70.4% | 1.2% |
| **HC-Pro** | 88.3 | 0.0% | 89.2% | 0.0% | 66.7% | 2.4% |
| **Nlb** | 87.7 | 0.5% | 88.4% | 1.2% | 59.8% | 3.1% |

#### Structural Alignment of PRSV-PK-CP

The Coat protein structures of three American isolates (Hawaii_S46722, Mexico_AJ012650, Colombia_KT275938), three South Asian isolates (Malaysia_AB044342, Thailand_AYO10721, Taiwan_X97251) and three closely related isolates (Meghalaya_MF356497, West Bengal_LC482263, Bangladesh BD2_MH397222), modeled through i-TASSER. The RMSD values of the all the aligned CP structures are mentioned in Table 7.

**Table 7.** PRSV-PK Coat protein structural alignment with the CP structure of the representative isolates

| **Representative clade** | **Structure alignment of PRSV coat protein (CP)** | **RMSD Score** |
| --- | --- | --- |
| **American** | PK-CP and Hawaii-CP | 0.603 |
|  | PK-CP and Mexico-CP | 0.719 |
|  | PK-CP and Colombia-CP | 0.812 |
| **South Asian** | PK-CP and Malaysia-CP | 0.776 |
|  | PK-CP and Thailand-CP | 0.570 |
|  | PK-CP and Taiwan-CP | 0.644 |
| **Closely related** | PK-CP and Meghalaya-India CP | 0.534 |
|  | PK-CP and West Bengal-CP | 0.577 |
|  | PK-CP and Bangladesh-CP | 0.656 |

#### Pakistan CP vs American CP

The .pdb file of the structural models of three American isolates were superimposed with the PRSV-PK CP structural model and the highlighted differences in the PRSV-PK amino acid residues are mentioned in the Figure 3.

**Figure Error!.**
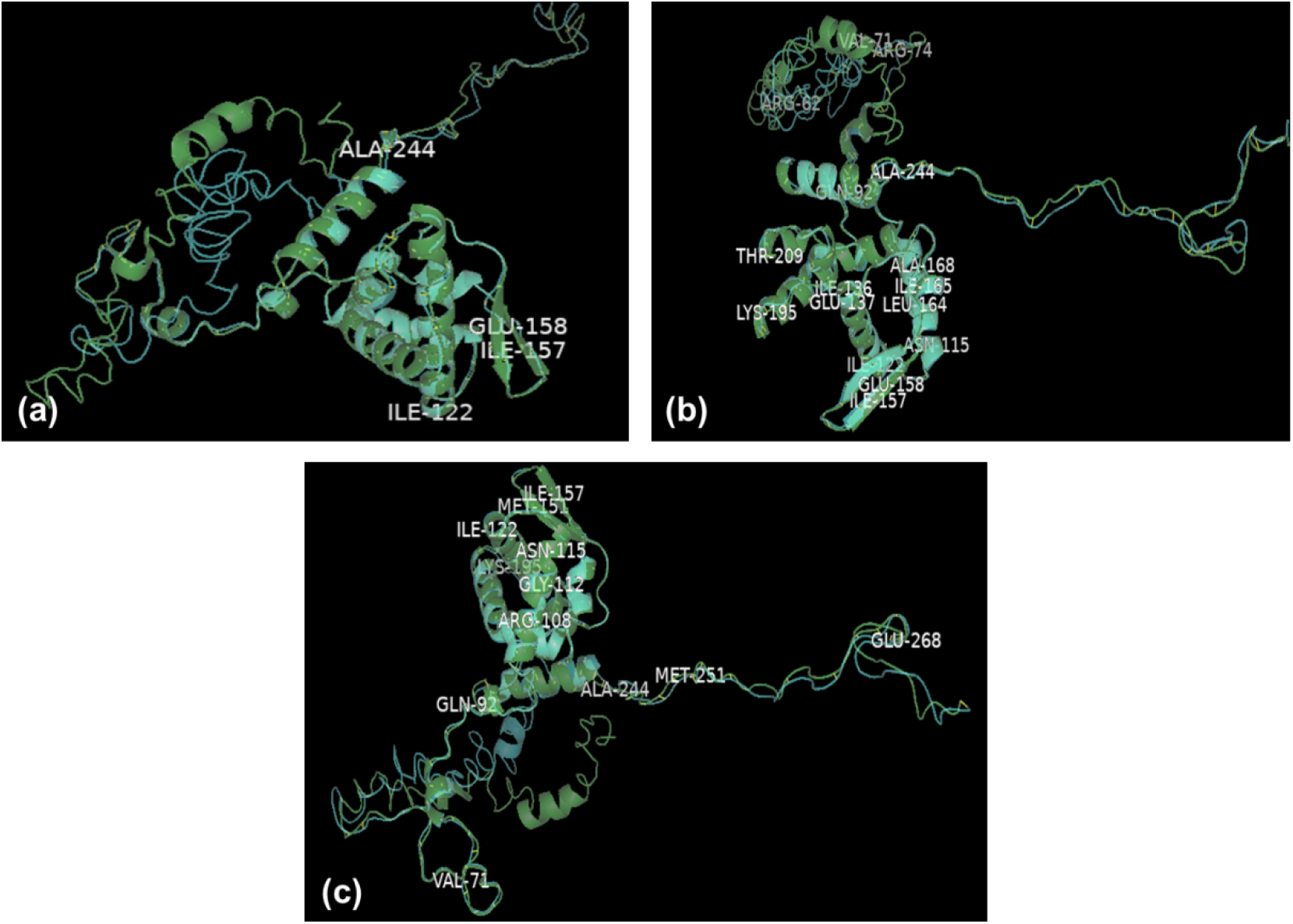
No text of specified style in document. Protein structure alignment via PYMOL (a) protein model superimposition of PRSV-CP from Hawaii and PRSV-PK-CP (**b)** protein model superimposition of PRSV-CP from Mexico and PRSV-PK (**c)** protein model superimposition of PRSV-CP from Colombia and PRSV-PK-CP. Structure in green color represents the structure of PRSV-PK-CP and structure in cyan color represents the structure of (a) Hawaii (b) Mexico (c) Colombia

#### Pakistan CP vs South Asian CP

The .pdb file of the structural models of South Asian CP were superimposed with the PRSV-PK CP structural model and the highlighted differences in the PRSV-PK amino acid residues are mentioned in the Figure 4.

**Figure 4.**
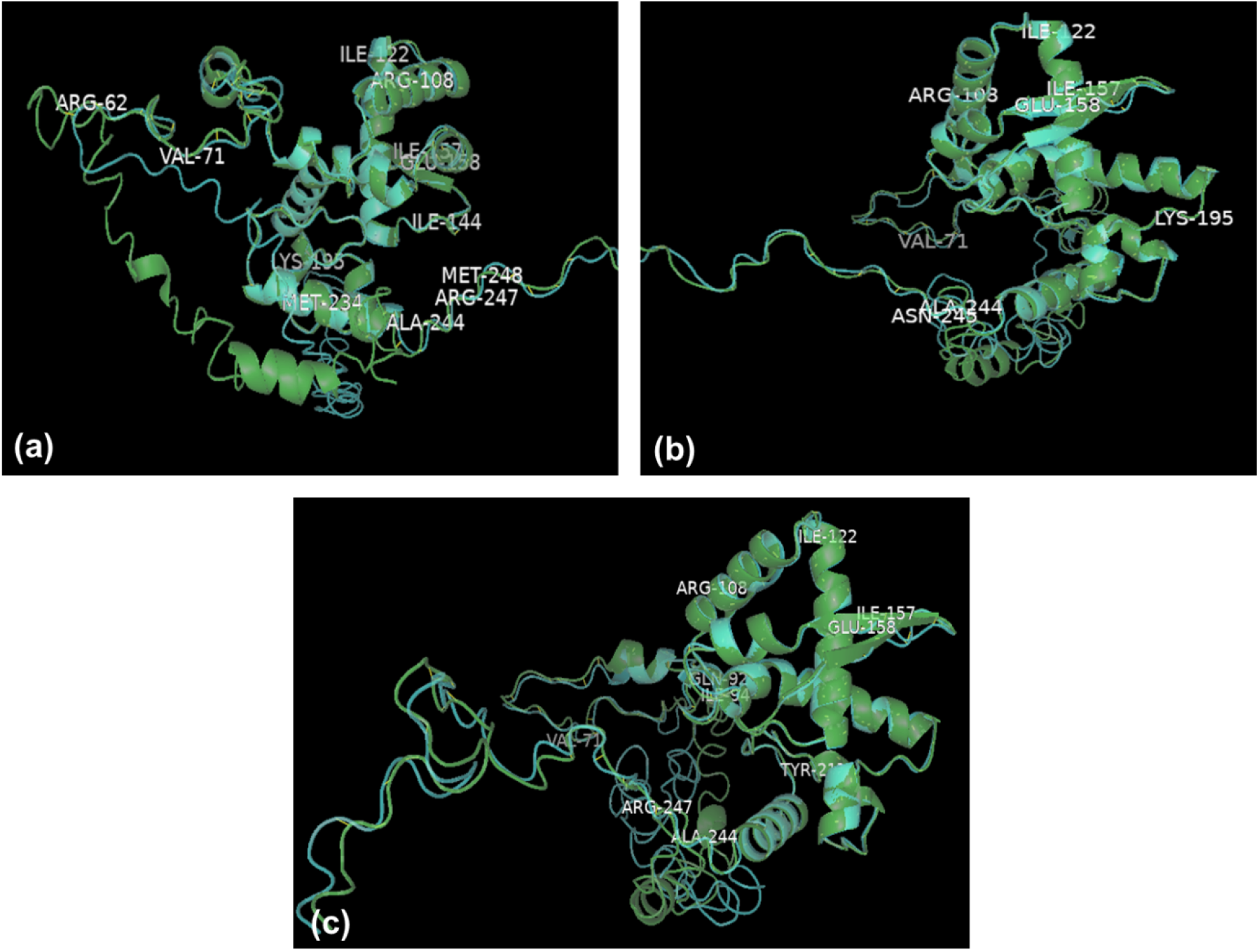
Protein structure alignment via PYMOL (a) protein model superimposition of PRSV-CP from Malaysia and PRSV-PK-CP (b) protein model superimposition of PRSV-CP from Thailand and PRSV-PK (c) protein model superimposition of PRSV-CP from Taiwan and PRSV-PK-CP. Structure in green color represents the structure of PRSV-PK-CP and structure in cyan color represents the structure of (a) Malaysia (b) Thailand (c) Taiwan.

#### Pakistan PRSV CP vs CP of closely related isolates

The .pdb file of the structural models of South Asian CP were superimposed with the PRSV-PK CP structural model and the highlighted differences in the PRSV-PK amino acid residues are mentioned in the Figure 5.

**Figure 5.**
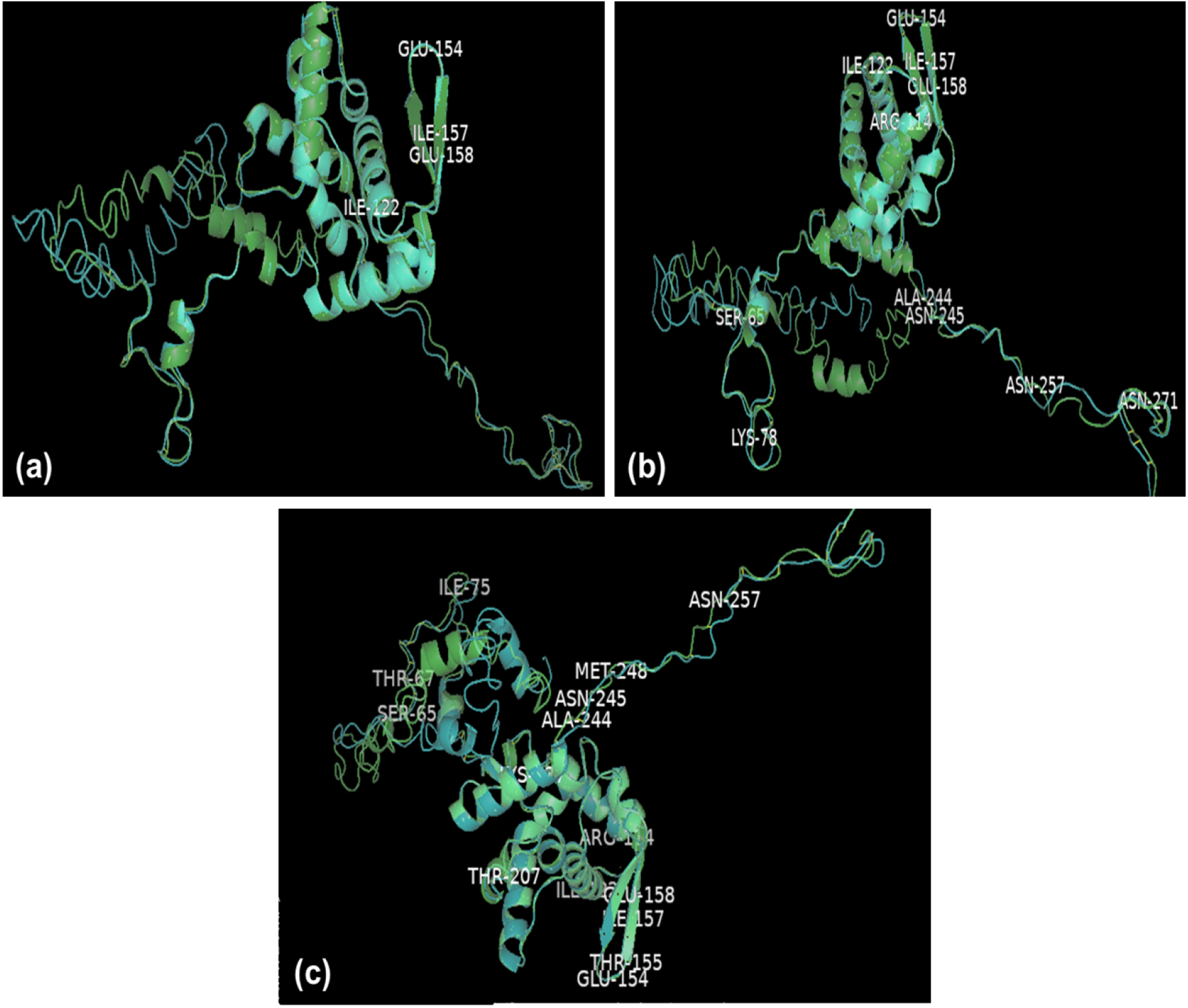
Protein structure alignment using PYMOL modeling via i-TASSER (a) protein model superimposition of PRSV-CP from Meghalaya-India and PRSV-PK-CP (b) protein model superimposition of PRSV-CP from West Bengal-India and PRSV-PK (c) protein model superimposition of PRSV-CP from Bangladesh and PRSV-PK-CP. Structure in green color represents the structure of PRSV-PK-CP and structure in cyan color represents the structure of (a) Meghalaya (b) West Bengal (c) Bangladesh

#### Docking Analysis

The PRSV-PK Coat protein (CP), HC-Pro and NIb interaction with host proteins was performed using ClusPro (Protein-Protein docking server). The lowest energy value for each interaction has been shown (Table 8).

**Table 8.**
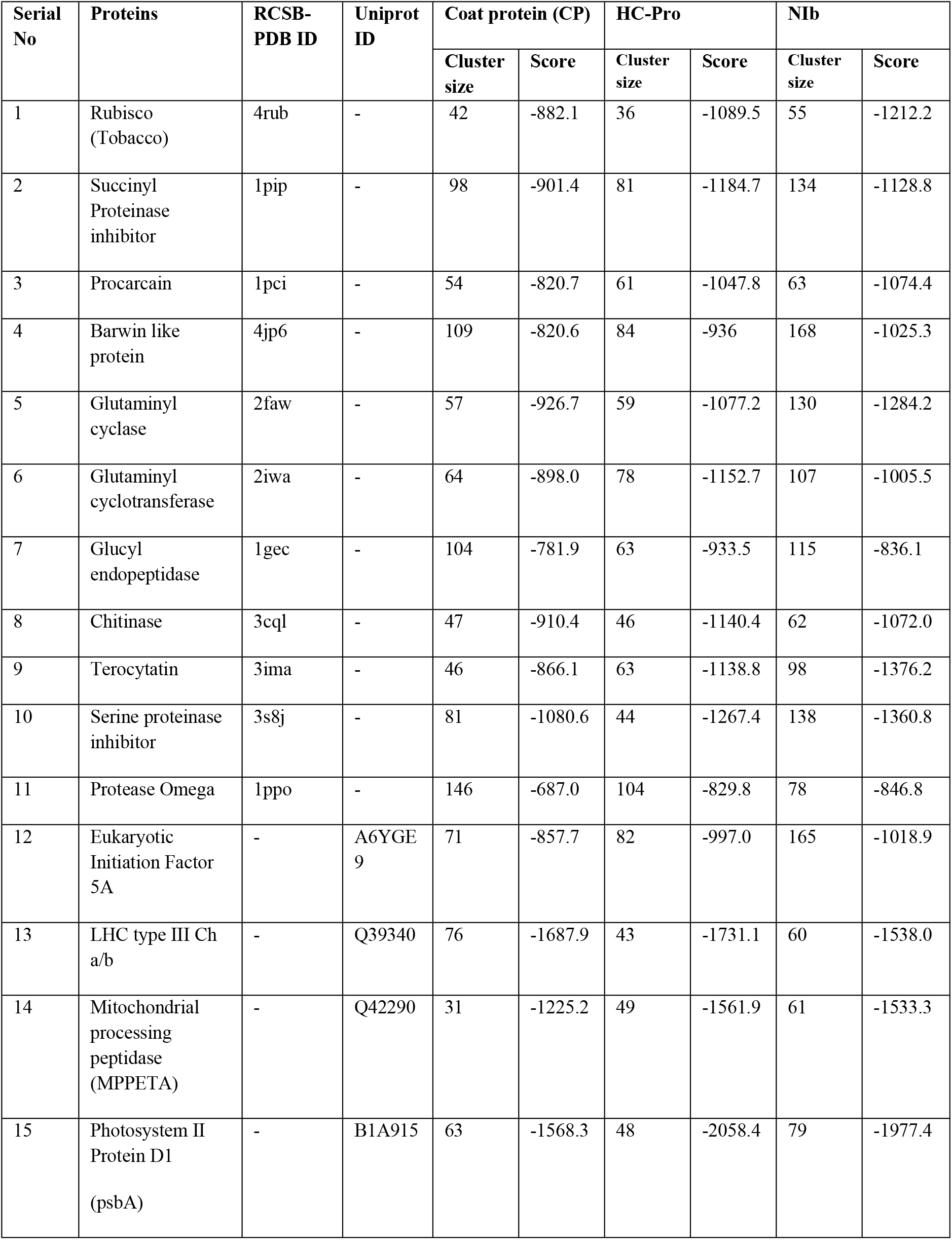
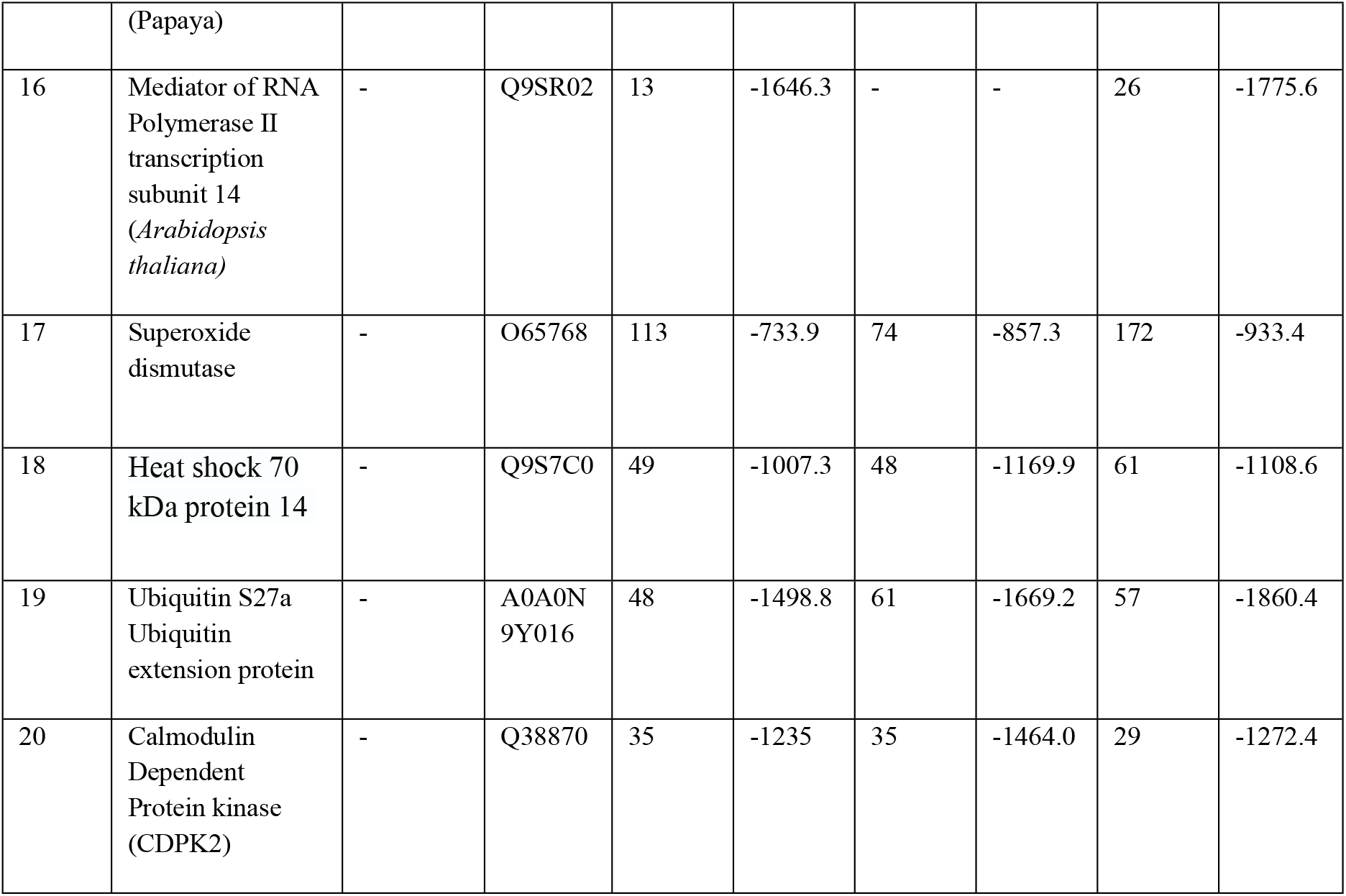
Docking of PRSV-PK CP, HC-Pro and NIb with the host proteins using ClusPro (Protein-Protein Docking Server)

## Discussion

Plant viruses hijack their host cells and direct host’s cellular pathways in provision of their infection cycle. For completion of replication, infection and systemic movement, they need to neutralize the complex host-defense mechanisms to ensure high vulnerability of the host [30]. Therefore, a deeper understanding of viral approaches which they employed for cell function manipulation as well as facilitation of their infection cycle is essential to produce stably resistant plant lines [31]. Structural bioinformatics is concerned with the computational approaches for protein structure and functions prediction. These structures provide a basis for functional analysis of experimentally derived structures, understanding the protein structure, number and types of epitopes, immunogenic portions, and suitability for antibody production, taxonomic studies, evaluation studies and virus diagnostics. In the current study three functionally vital proteins of PRSV-PK strain has been modeled using compatible modeling servers. The nucleotide based genetic diversity studies on CP gene of PRSV-PK strain has already been conducted extensively [6,9]. However, the diversity exploration of virus genome is not only confined to molecular approaches including recombination and phylogenetics, rather structural information of virus proteins has shown significant contribution towards virus diversity in terms of its structure-function relationship [32]. Henceforth, the underlying study has been conducted realizing the niche regarding the structural knowledge of essentially important proteins of PRSV. The modeled structures were then subjected to functional mapping which provided the valuable insight into their probable interactive role within the host.

The physiochemical properties of CP, HC-Pro and NIb determined using Expasy’s ProtParam server [20], allows the computation of various physical and chemical parameters for a given protein. Grand average hydropathy (GRAVY) values of all the amino acids divided by the number of residues in the sequence, for the CP, HC-Pro and NIb protein in our study was -0.842, -0.435 and -0.328 respectively. The negative GRAVY values of the given proteins indicate their better interaction with water (see Table. 4). The physicochemical characterization of several other proteins have also been investigated previously [22,23,33,34]. The secondary structure prediction of the essential PRSV proteins was performed using a newly discovered method known to be Self-optimized prediction method (SOPMA) which was reported previously with improved success rate in secondary structure prediction [34]. The results revealed the domination of random coil in case of CP protein structure while Alpha helix dominated among HC-Pro and NIb secondary structure elements. Similar prediction method was employed by [23] for CP sequence of MYMIV and the domination of alpha helix was also predicted by [22]. Structural modeling along with biochemical analysis and physicochemical characterization are vital source to map the conserved motif and hyper variable area associated with N-terminal of CP Structural modeling [32]. The PRSV-PK proteins modeling were performed by utilizing most recent homology modeling tools including Swiss-Model, Phyre2 and i-TASSER. Phyre2 built model showed 100% confidence for 67%, 63% and 20% sequence identity of residues for PRSV CP, HC-Pro and NIb proteins (Table 3). The Swiss-Model showed sequence identity of 63.40%, 62.42% and 16.49% for CP, HC-Pro and NIb genes respectively (Table 4). The sequence identity values of Phyre2 and Swiss-Model are more or less closely related with highest sequence identity value for CP whereas the lowest identity for NIb. The results from the models predicted by i-TASSER indicated residue coverage of 67% for CP, 63% for HC-Pro and 20% for NIb respectively (Table 5). The structure of CP, HC-Pro and NIb modeled by each modeling server is shown in Figure 2. Interestingly the genomic structure of flexous filamentous viruses including those of *Potyviridae*, family, differs significantly, especially their CPs exhibit least similarity on the basis of sequence, however over all architecture of their virions is quite similar at least at a low resolution [35]. Henceforth, suggesting common homologous folds is shared by CPs from flexous rod-shaped viruses [36]. Similar common folds were observed in the overall architecture of modeled CP from different region.

Model quality evaluation is critical in homology modeling to ensure that model’s stereochemistry is consistent with typical values of crystal structure. The model’s quality estimated by Ramachandran Plots using PROCHECK tool shows that the maximum portion of the models lies in the ‘allowed’ region and less than 0.5% lies in ‘disallowed’ region which indicates that the good efficacy of the models. Ideally, over 90% of residues in the core region represent the good quality structure. Although in this study the percentage residues within the allowed regions were not above 90% but the percentage of the residue of all the modeled structures of the study lies in the allowed region was strikingly greater than those lies in disallowed region. Further, the results also showed the phi-psi torsion angles for all the structural residues with the exception of residues at chain termination.

The alignment of amino acid sequences of CP from different inter geographical regions using Constraint-based Multiple Alignment Tool (COBALT-NCBI) predicted the intrinsically disordered regions towards the N-terminal region presenting high sequence variation and high flexibility. These finding complies with the reports suggesting high polymorphism in the N-terminal region associated with functional versatility leads to multiple interactions [37]. Alignment of CP modeled structures of PK strain with CP structures from different geographical locations showed the N-terminal region being highly flexible. Further, the structure alignment of PK-PRSV-CP with CP structures from American clade indicated the amino acid substitution was high at the core region compared to C-terminal which may influence the virion assembly (Figure 3). More residual differences were observed between the aligned structures of PRSV-PK-CP and PRSV-CP-Colombia showing the highest RMSD value of 0.812 (Table 7). However, the overall residual differences do not affect the structural topology of these proteins. The same observations were noted among the alignment of PRSV-CP structure and CP structures from the isolates of South Asia (Malaysia, Thailand and Taiwan) (Figure 4) and those of closely related including India and Bangladesh (Figure 5). The minimum root means square deviation (RMSD) values of 0.534 and 0.5377 were observed between the atomic residues of PRSV-PK CP with Meghalaya-CP and Bangladesh-CP (Table 7), which may suggest that closely related structures share a common protein fold as well as functional roles.

The preliminary *in-silico* structural prediction of potyviral HC-Pro was remained unsuccessful. However, the structural confirmations the protein adopts showed that even its well characterized protease domain was restricted to the C-terminal protein region [17] thereby hindered its substantial prediction. Further the oligomerization ability of HC-Pro [38] has also been attributed towards unsuccessful domain interaction studies. Later, several studies explored the structural knowledge of HC-Pro with a functional correlation, particularly impact of its oligomerization on aphid transmission [39]. NIb, another functionally important gene of PRSV that codes for the RNA-dependent-RNA polymerase responsible for RNA replication [40]. The structural arrangement of viral RdRps have revealed their classical closed right hand architecture consisting of three subdomains including palm, thumb and fingers [19].

Once the structural studies will be accomplished the functional aspects of the genes and association with other molecules will be more comprehensible. Therefore the analysis of the protein structures of these genes further aids in exploration of their flexible association leading to vector transmission [32]. The hypothetical interlinking of functionally active proteins of PRSV-PK strain and with each other and with molecules within the invaded host cell has been depicted (Fig 6). CP gene of PRSV is responsible for various functions including virion formation, cell to cell movement and systemic movement [41]. The terminal region of CP contains a conserved motif ‘DAG’ that interacts with HC-Pro to ensure aphid transmission of the virus [42,43]. The genetic compatibility between CP and aphid transmission motif has been found to be critical for functional maintenance in genetically distant host vectors [44].

**Figure 6.**
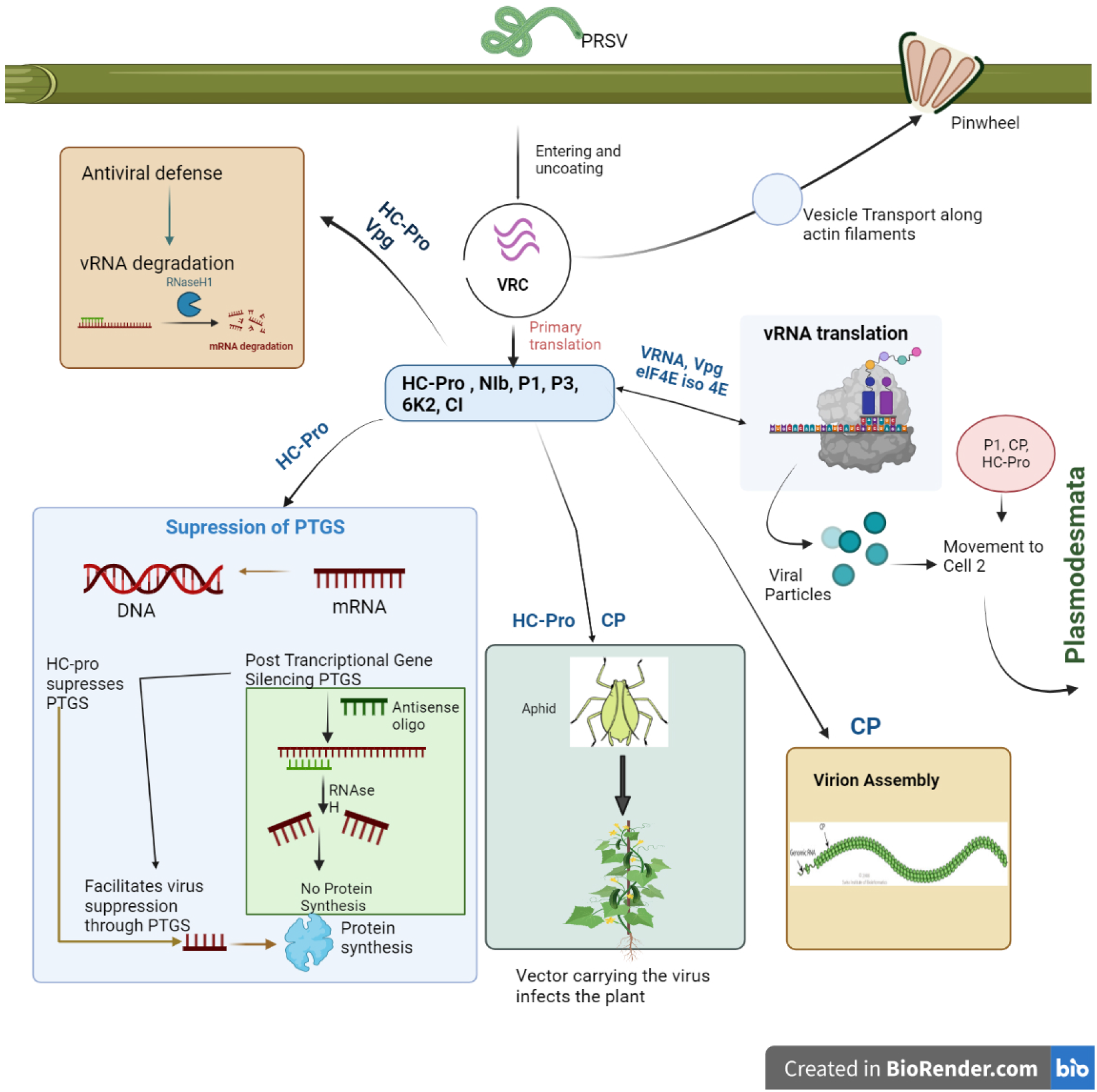
Functional interactions of PRSV proteins inside an infected plant cell. CI, NIa, NIb, HC-Pro and P3 plays interacts with host cellular machinery, HC-Pro along with the CP gene aids in vector transmission of PRSV to other plants. HC-Pro also acts as a strong silencing suppressor of Post transcriptional gene silencing (PTGS). VPg domain of NIa-Pro interacts with host translation initiation factors eIF4E and helps in virus RNA translation thus aids the virus in causing infection. P1, HC-Pro and CP play a role in cell-to-cell movement of PRSV.

The typical structure of PRSV-PK-CP coat protein depicted that it harbors a conserved DAG motif (Asp-Ala-Gly) [42]. The presence of DAG motif was somewhat consistent in all the aligned protein sequences thus signifying the presence and involvement of the motif in aphid transmission. The amino acids in the vicinity of DAG motifs are also equally contributing in the virus transmission [42]. However, it has been reported that the DAG motif is not universally conserved and there is not only a variation in amino acid sequence but also in location of the DAG motif in several other potyviruses with frequent replacement of DAG with NAG, NVG or DTG [32]. Similarly *Bean yellow mosaic virus* (BYMV) contains a NAG or a KAG motif instead of DAG motif. Our results report that the CP of isolates *viz*, Meghalaya and Taiwan lacks the typical DAG motif. The CP of the Taiwan isolate possess DTG motif instead of DAG [32,44] (Figure 7a). Apart from DAG motif the other conserved motifs in CP, NAG, WCIEN and QMKAAL [45], have been mapped on PRSV CP structure (Fig 7b). The core domain has been reported to be critical for RNA encapsidation and virion formation [40,46]. Henceforth, mutational alterations in specific parts may leads to differential changes in virus properties [37]. As the high diversity in N-terminal region of CVYV and Cassava brown streak viruses seem to be determinant of whitefly transmission, however the deletion of 13aa in N-terminus would leads to the loss of transmission [37].

**Figure 7.**
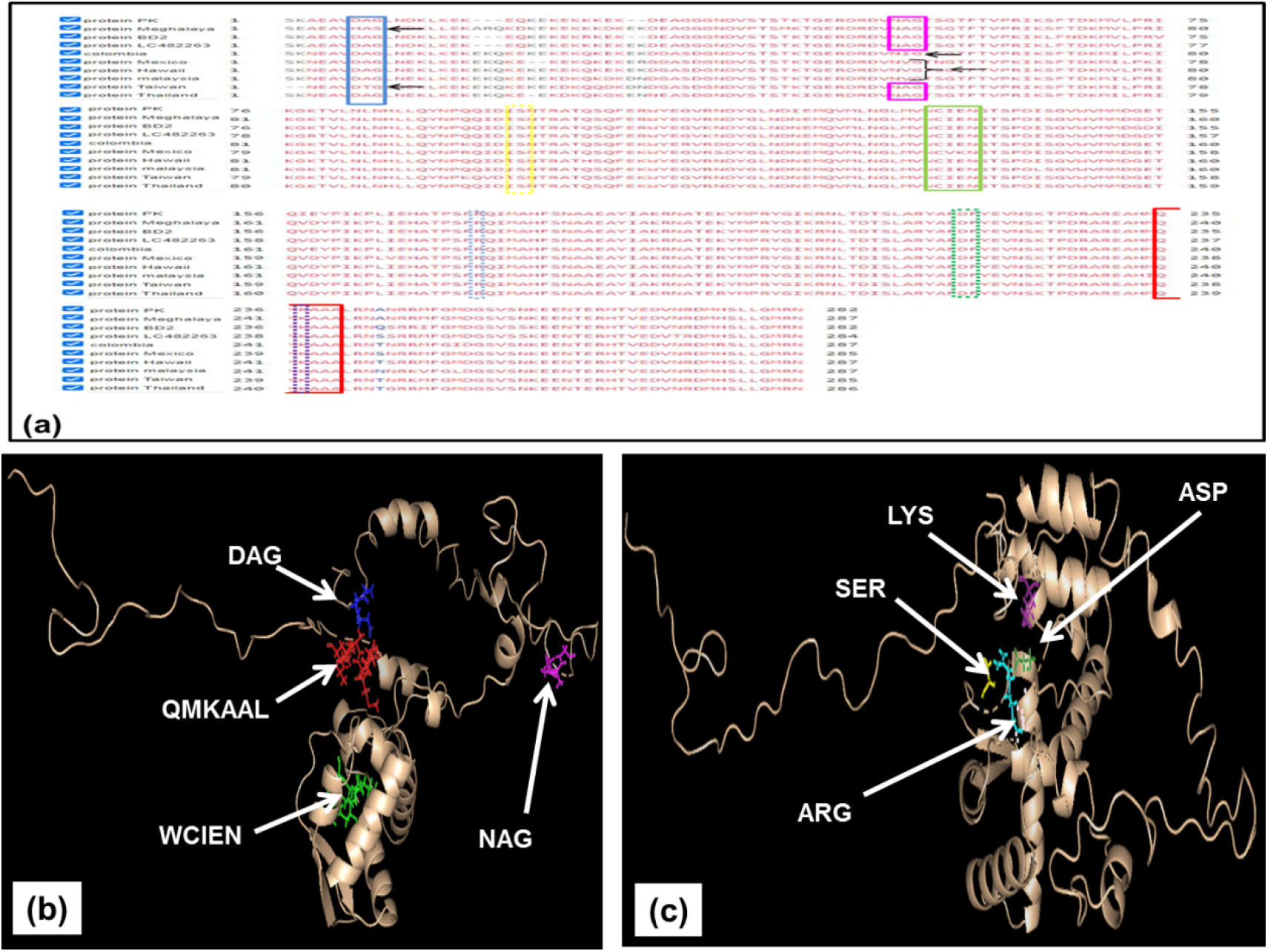
PRSV-PK-CP showing conserved motifs and critical amino acids (a) Amino acid alignment of Coat protein. The motifs inside the colored shapes are conserved and the black arrows indicate the mutation across the conserved motifs among CP of PRSV from different geographical locations (b) Ribbon modeled PRSV-PK-CP structure showing conserved amino acids, the conserved motifs are shown as colored lines each color corresponds to the colored assigned to conserved motif in amino acid alignment alignment, green: WCIEN motif in the core domain of CP, Red: QMKAAL motif at the C-terminal domain of CP, Dark blue : DAG motif at the N-terminal region, Magenta: NAG motif at the N-terminal region (c) Critical amino acids which are found to interact with ssRNA and aids in virion formation are conserved among the PRSV-CP isolates from different geographical locations. Yellow: Serine, Cyan Blue: Arginine, Forest green: Aspartic acid, Deep Purple: Lysine.

Similarly the interactions of *Tomato mosaic virus* (ToMV) with the host factor promotes the viral long distance movement in the tobacco plant [47]. CP of *Potato virus* X (PVX) interacts with the ER proteins of *N. benthamiana* which assisted viral replication in host cells [48]. CP protein of *Rice dwarf virus* (RDV) hinders the auxin signaling by targeting the OslAA10 protein of rice thereby resulted in enhanced severity of viral infection [49]. The two homologous protein of *N. benthamiana* interacts with the CP gene of invading PVX virus and interfere in its encapsidation and movement [50]. Another report revealed that *Cucumber mosaic virus*-cholrosis inducing strain interacts with choloroplastic ferredoxin I (Fd I) protein, whereas the green mosaic-inducing strain of CMV did not showed this interaction, which suggested that the protein interactions vary with the type of strain of the same virus [48]. The critical amino acids which are known to conserve in several potyviruses i.e *Pepper mosaic virus* and *Water melon mosaic virus* are potentially involved in interaction and binding with ssRNA [46]. Consistent with these findings the four critical amino acids which are serving as RNA binding sites have been mapped on the CP structure of PRSV-PK strain (Fig 7c). These conserved RNA binding residues in CP’s of flexous filamentous viruses provide a clear target for antiviral compounds that may interfere with the virion assembly of many economically significant plant viruses [46].

Functional motif identification in HC-Pro suggested the division into three main regions, an N-terminal cysteine rich region metal binding motif has been known to be responsible for symptoms severity, systemic infectivity and aphid transmission [16]. The conserved motif KITC at the N-terminal domain has been known to conserve in all aphid transmitted potyviruses, with functional interaction with aphid stylet [51]. The core region is comprised of two RNA binding domains, as evident from high lysine, arginine and asparagine content present in this region. Probably, the former domain having the conserved motifs including FRNK, WG and CDNQLD has been known to play a role in genome amplification [13]. Similar conserved motifs have been mapped from PRSV-PK-HC-Pro (Fig 8). While, the later domain with the conserved motifs including YHAKRFF, GY, PNG and AIG have been known to play part in PTGS suppression, while other conserved motifs are associated with systemic virus movement and synergism [51]. Similar functional associations of conserved motifs of HC-pro core domain has been described for *Banana bract mosaic virus* (BBrMV) [13]. Consistent with these findings, similar conserved motifs have been mapped in the core domain of PRSV-PK-HC-Pro and in associated PRSV-HC-Pro from other geographical regions (Fig 8). The C-terminal domain has shown to harbor proteinase activity and plays a crucial role in cell to cell movement. The conserved motifs of this domain NIFLAML, AELPRILVDH, LKANTV and VG as described by [51] has also been mapped in PRSV-PK-HC-Pro. Another important functional motif i.e PTK has been found to be evolutionary conserved in all potyviruses and has its probable role in binding of HC-Pro to viral CP. Henceforth, the presence of many conserved motifs within this region confirms the fundamental role of this domain as proteolytic enzyme in all the potyviruses irrespective of their host. The amino acid alignment of PRSV-PK-HC-Pro provided an insight into the highly conserved functional motifs in all three domains (Fig 8). The insight into the functional motifs of HC-Pro therefore suggested that its highly conserved domains are primarily involved in vital functions of post transcriptional gene silencing (PTGS) suppression and proteolytic activity which are pivotal in plant-virus cycle. The structural interface and conserved motif provided an insight into the interaction between viral-host proteins [51].

**Figure 8.**
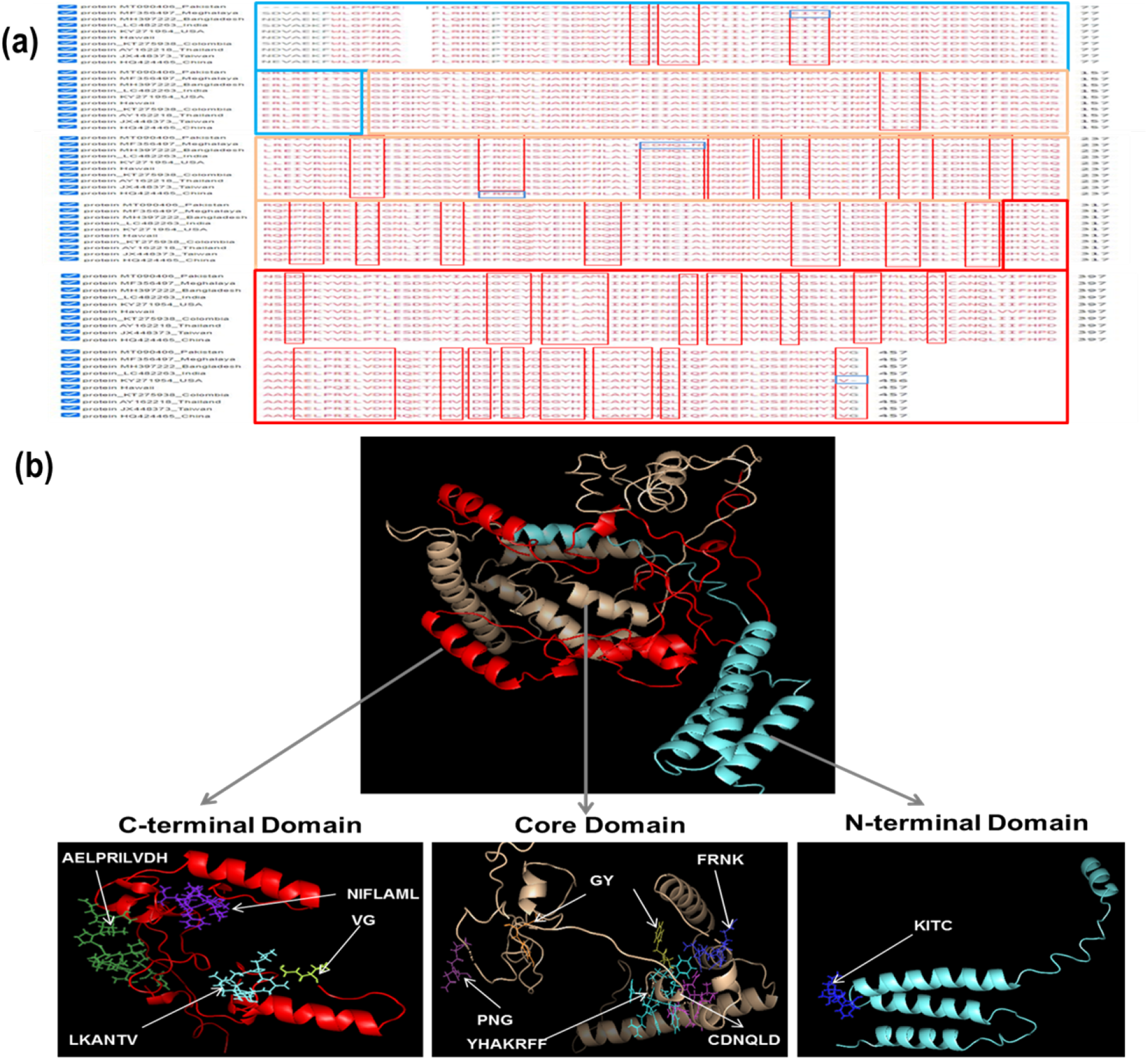
The amino acids sequences of Helper Component Proteinase (HC-Pro) of (a) Aligned protein sequences of *Papaya ringspot virus* representative isolates (b) the color of three domains corresponds with the color of three big squares on the protein sequence alignment. The colored line structure indicates the motifs in each domain. All the conserved motifs within each domain are shown in red boundary. C-Terminal domain (Red), Core domain (selmon), N-terminal domain (Cyan)

Another functional viral protein NIb, the viral RdRps have been known to be structurally arranged in a classical closed right hand architecture consisting of three subdomains including palm, thumb and fingers. Most conserved viral RdRps motifs are found to be in palm subdomain, whereas the motifs within fingers and thumb subdomains are found to be diverse. The palm subdomain of PRSV-PK NIb possess five structurally conserve motifs including CVDDFN, CDADGS, GNNSGQPSTVVDNTLMVL, GDD and ALIE(K)SWG as described [52]. The GDD motif has been known to be fundamental for RdRp activity. The finger domains possesses two conserved motifs [53]. Similar conserved motifs were identified in finger domain of PRSV-PK-NIb protein. It has been suggested that potyviral NIb exhibits relatedness to animal picornaviruses [36,54], therefore may follow the same regulatory pathways to that of poliovirus RdRp. The seven conserved motifs including SLKAEL, CVDDFN, CHADGS, GDD, [A/S]M[I/V]E[S/A]WG, FTAAP[L/I][D/E], and GNNSGQPSTVVDNTLMV have been consistently reported in potyviral NIbs [55,56] and have been identified in PRSV-PK-NIb (Fig 9). The functionally conserved motifs of NIb protein of PRSV-PK and the associated PRSV isolates from geographical regions (Distant, Related) have been mapped on the aligned protein sequences (Fig 9). Variation in NIb confines to the C terminal part near the junction with the CP. However, variation at the NIb-CP junction might be related to efficiency of polyprotein processing by NIa-Pro.

**Figure 9.**
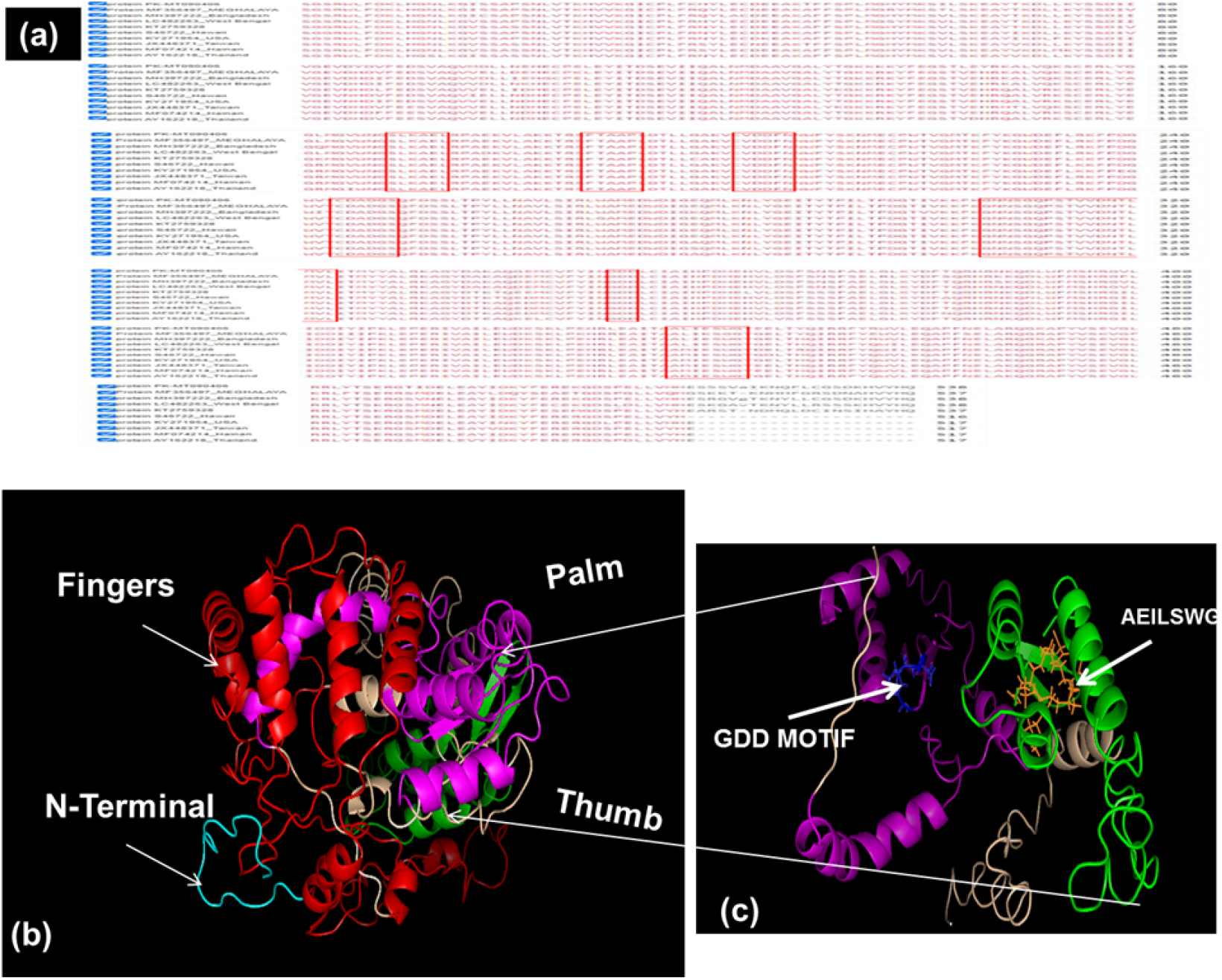
The amino acids sequence alignment of Nuclear Inclusion Protein b (NIb) of *Papaya ringspot virus* representative isolates, structural representation and critical domains. (a) Aligned protein sequence of PRSV-PK-NIb with the NIb of representative PRSV isolates (b) Structural domains of NIb, Red: Fingers, Pink, Palm, Green: Thumb: Cyan: N-terminal. (c) conserved motifs are shown in Colored lines within the finger and Thumb domain. Multiple alignments were performed with Constraint-based Multiple Alignment Tool (COBALT-NCBI).

Viral protein interactions with the host protein lead to successful replication, pathogenesis and infection of PRSV. Hence, to understand these mechanisms a computational analysis of PRSV-PK-CP with essential host proteins was carried out in this study. The *in silico* protein docking analysis predicted the interactions of invading PRSV-CP, HC-pro and NIb with distinct host proteins, functional switching of these protein’s with those of the host which leads to successful invasion, replication and propagation of virus. Viruses can mainly manipulate chloroplast and along with that they also alter other organelles including nucleus, endoplasmic reticulum (ER), mitochondria, and peroxisomes [31]. Thus higher insight into the interaction would reveal the possible pathways through which the virus infection cycles could be turned off during possible phase of its infection cycle. Coat protein of RNA viruses holds many other activities a part from their well-recognized structural activity [57]. Our study of Coat protein interaction with the host revealed the good interactions with LHC type III chlorophyll a/b binding protein, PsbA, Photosystem II (PSII) is a light-driven water: plastoquinone oxidoreductase (Fig 10) that uses light energy to abstract electrons from H_2_O, generating O_2_ and a proton gradient subsequently used for ATP formation, Mediator of RNA polymerase II transcription subunit 14 involved in transcription regulation and signal transduction within the host and mitochondrial peptidases which are known to play protein metabolism within the host cell. Previously, *in vitro* approaches that determined the interaction of CP inside papaya identified, twenty-three host proteins interacted with invaded viral CP. Those proteins were mainly involved in cellular metabolism, transcription, translation, carbohydrate metabolism, protein metabolism, stress response, photosynthesis, nucleotide metabolism, respiration and lipid metabolism [58]. The interaction values in our study has provided the probable functions of these proteins are within the host further suggested whether the interaction between host and virus though these proteins is compatible or not. However, here it can perceived that coat proteins recruits the host proteins in possibly two ways, that it either shuts off the host proteins to support its own functions or activates the host proteins which further acts against them resulting in host defense. In current study we can perceive that CP being the major symptom determinant interacts with the chloroplast or the proteins associated (as indicated by high energy values of interaction) with the chloroplasts including PSII complex as reported by [59], or with LHC and psbA resulting in the impaired function of chloroplast which eventually came up with mosaic and chlorosis symptoms on plants. Chloroplast plays an important role in plant defense through production of Reactive Oxygen Species (ROS) which can directly prevent infection through stimulation of programmed cell death or indirectly by activating defense related genes in nucleus [60]. Thus, virus must suppress the chloroplast mediated defense in any way to ensure its replication. Further, the virus interaction with the host chloroplast through protein-protein interaction is of utmost significance [61,62] as symptoms development is the first indication of disease onset. However, the mode of interaction can be direct or indirect, as several other reports suggested that potyviral CP interactions with the host protein resulted in the formation of membrane vesicles which in turn improve the viral replication and developed vesicle shuts off the chloroplast functions [63,64]. Thus interactions between CP and host proteins are thus crucial for progressive infection by the invading virus [65]. Apart from recruitment of host proteins by CP in its own favor, CP also interacts with invaded papaya protein resulting in host defense activation against the virus. There exist several examples where the heat shock protein 70 (HSP70) along with its co-chaperone CPIP governs the potyviral CP which in turn inhibit replication [57]. Similarly, in our study the high interaction of PRSV-PK CP with Heat shock protein (HSP70) (Fig 10) is also an indication of the association between the two which would be much likely in favor of host. Further, *in-silico* analysis docking based interaction study revealed that CP dimers of *Sesbania mosaic virus* interact with three potential candidate proteins of host during initial phase of infection. Most importantly these proteins belong to lectins and canavalin families and are involved in antiviral defense in plants [66]. In our study docking analysis of PRSV-PK-CP also showed high interactions with RNA polymerase II transcription mediator protein of host which is responsible for transcriptional regulation. It is assumed that upon the entry, viral CP interacts with RNA Pol II Transcription mediator which gets activated in response to stress and induce a cascade of abscisic acid signaling pathways and activates host’s defense. Thus this association may leads to incompatible virus-host interaction resulting in host defense.

**Figure 10.**
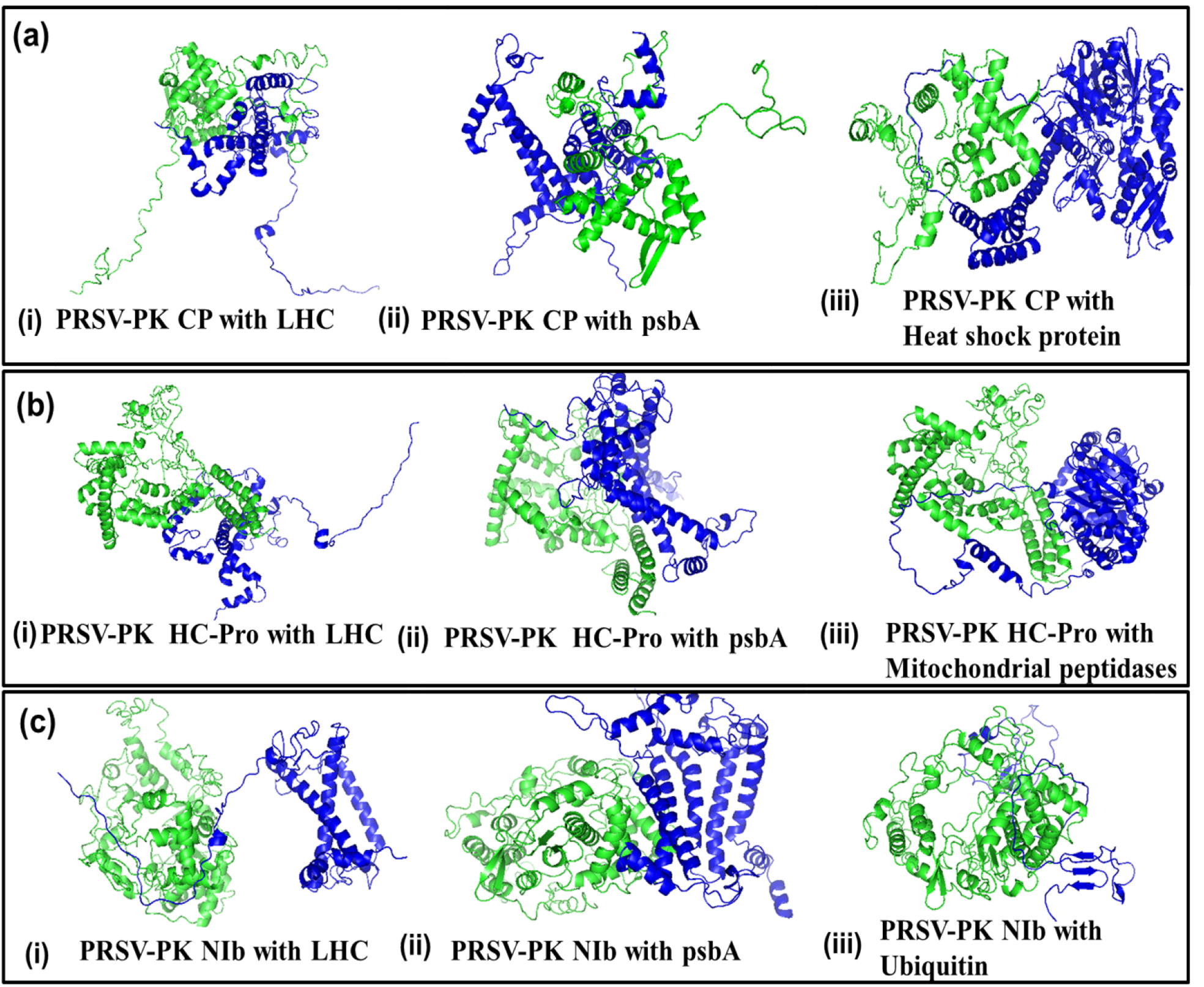
Protein-Protein Docking analysis of *Papaya ringspot virus* proteins CP, HC-Pro and NIb with host proteins using ClusPro Server, (a) Shows the docking analysis of PRSV-PK -CP with (i) LHC, (ii) psbA (iii) Heat shock protein. (b) Shows the docking analysis of PRSV-PK-HC-Pro with (i) LHC, (ii) psbA (iii) Mitochondrial peptidases, (b) Shows the docking analysis of PRSV-PK-NIb with (i) LHC, (ii) psbA (iii) Ubiquitin. The green Colored protein indicates the PRSV-PK proteins and Blue colored proteins are the host proteins.

Several studies reported the interaction of HC-Pro with translation initiation factor which augment virus replication [67,68]. In our study the viable interactions of HC-Pro with the host Light harvesting complex (high interaction values mentioned in Table 8 points out that it also accumulates at thylakoid membrane but here it intensify the symptom, as CP is already there for chlorophyll destruction so it may work in proximity with CP and leads to more severe symptoms through further disruption of chlorophyll function. There is also a probability that HC-Pro, of other potyviruses infecting papaya follow the same route ad through compatible interactions with the host Light harvesting complex they synergistically act with each other which lead to symptoms severity. Further the high interaction of HC-Pro with Ubiquitin extension protein (Table 8) proposed that it might leads to unfavorable interactions for the virus, as ubiquitin-mediated 26S proteasome has been reported previously to inhibit viral infection in plants [69]. However further insight through *in-vitro* approach is needed to confirm the exact pathway that Ubiquitin might adopt to inhibit silencing suppressor activity of HC-Pro.

The interactions of viral proteins with nuclear associated proteins provide more relevance to viral movement within the host than its replication. Nevertheless, the destabilization of nucleus by virus turn off the RNA silencing events of host thus enables the virus to surmount the adjacent cells and eventually the entire tissues [31]. However, being RNA viruses the nuclear recruitment is not possible directly but the virus can pass these limitations by producing signals through its protein that interacts with nuclear proteins to get themselves in potyviruses has been known to possess two nuclear inclusion proteins, NIa and NIb which interacts with nuclear host proteins and thereby facilitates the assembly of viral replication complex within the host nucleus [40].

Therefore, NIb has been known to be a key player in diverse virus host interactions by inducting the host proteins into viral replication complexes (VRCs). NIb possesses nuclear localization signals (NLSs) because of which it accumulates in the nucleus in the form of crystalline nuclear inclusions (NIs), mutational changes in NLSs interrupt its localization and terminates the genome replication. The strong interaction of PRSV-PK NIb with Ubiquitin S27a ubiquitin extension protein (Table 8) and Fig 10 which is involve in RNA binding and Translation, suggests that association of NIb by which it recruits the host translational machinery for its own usage.

The interaction of potyviral NIb with PABP2, PABP8, eukaryotic elongation factor 1A (eEF1A), heat shock cognate protein (Hsc or HSP) 70-3, and an RNA helicase-like protein in *Arabidopsis* (AtRH8) has also been reported. Hence the interaction of NIb with host proteins has been shown to be mandatory, to deliver the functional RdRps in VRC [19].

Papaya being eminent crop of South Asia also has wide spread cultivation in Pakistan. There are reports of wide spread cultivation of papaya in Sindh, Province [6,7]. Since 2013, the attack of *Papaya ringspot virus* leads to dramatic reduction in fruit quality, vigor and ultimately the overall production. The immediate solution to mitigate the virus has become essential to regulate papaya cultivation in order to enhance the country’s economy. Lack of adequate knowledge regarding structural parameters of essentially important proteins of the virus will compromise the functional exploration associated with the structures. Computational based study for physiochemical structure analysis will help in understanding the proteins structure and provide the chance to compare PRSV-PK essential protein sequence with other countries on visual basis. The present study would also be helpful for development of broader disease resistance/management strategy for the ring spot virus by discovering PRSV diversity based on protein structure and the functions they play within the host cell.

## Conclusion

Homology modelling is an imperative application of structural biology which narrows the gap between protein sequence and its experimentally determined structures. Nevertheless the structural prediction using different computational tools have provided a useful insight in structure-function relationship and interaction of viral proteins with the host via protein-protein interactions. The functional protein mapping along with protein-protein docking analysis showed the virus-host interaction that is much more important to understand the infectivity mechanism of virus and activation of cellular response and defence against viruses.

## Acknowledgements

We are greatly thankful to Higher Education Commission Pakistan (HEC) for funding the research work under NRPU project no. 3551.

